# Optogenetic activation of heterotrimeric Gi proteins by LOV2GIVe— a rationally engineered modular protein

**DOI:** 10.1101/2020.08.17.253781

**Authors:** Mikel Garcia-Marcos, Kshitij Parag-Sharma, Arthur Marivin, Marcin Maziarz, Alex Luebbers, Lien T. Nguyen

**Author notes:** Graduate Curriculum in Cell Biology and Physiology, Biological and Biomedical Sciences Program, University of North Carolina at Chapel Hill, Chapel Hill, North Carolina 27599, USA.

## Abstract

Heterotrimeric G-proteins are signal transducers that mediate the action of many natural extracellular stimuli as well as of many therapeutic agents. Non-invasive approaches to manipulate the activity of G-proteins with high precision are crucial to understand their regulation in space and time. Here, we engineered LOV2GIVe, a modular protein that allows the activation of Gi proteins with blue light. This optogenetic construct relies on a versatile design that differs from tools previously developed for similar purposes, i.e. metazoan opsins, which are light-activated GPCRs. To make LOV2GIVe, we fused a peptide derived from a non-GPCR protein that activates Gαi (but not Gαs, Gαq, or Gα12) to a small plant protein domain, such that light uncages the G-protein activating module. Targeting LOV2GIVe to cell membranes allowed for light-dependent activation of Gi proteins in different experimental systems. In summary, LOV2GIVe expands the armamentarium and versatility of tools available to manipulate heterotrimeric G-protein activity.

**GRAPHICAL SUMMARY:** 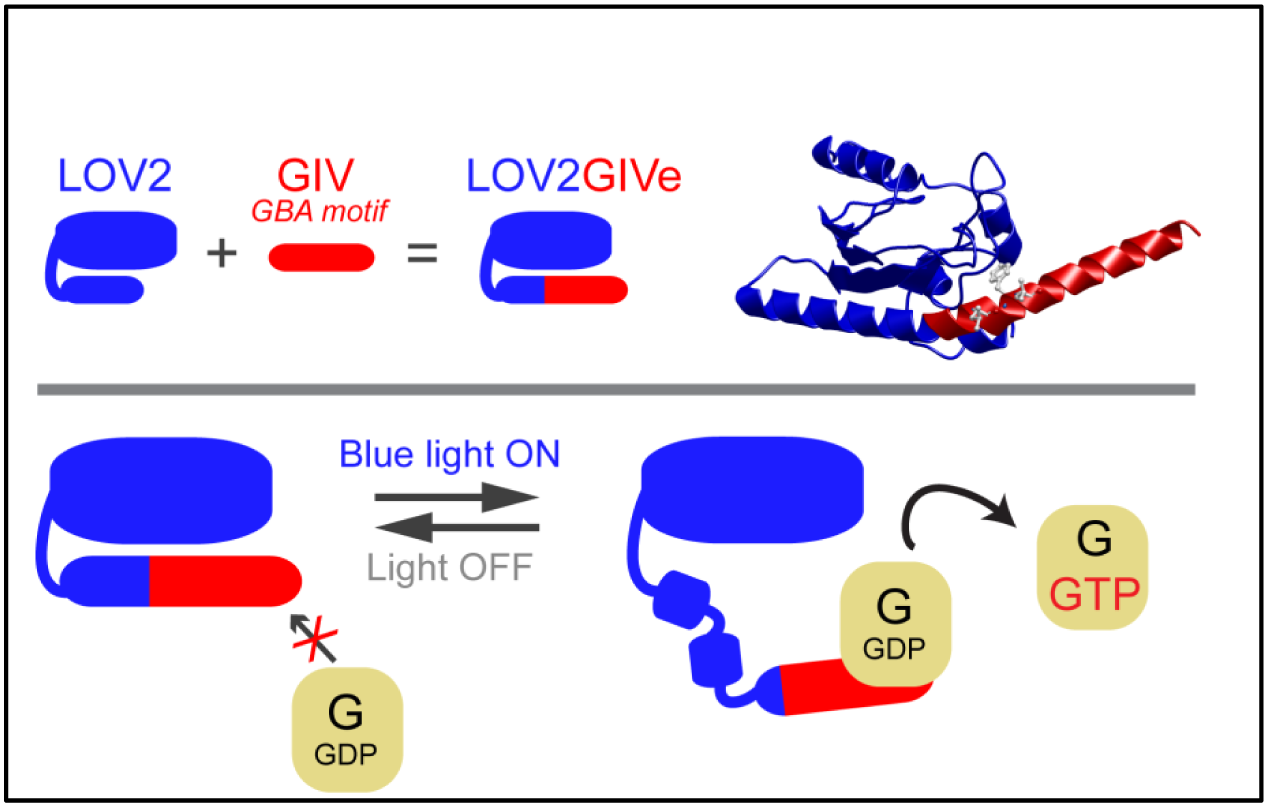

## INTRODUCTION

Heterotrimeric G-proteins are critical transducers of signaling triggered by a large family of G-protein-coupled receptors (GPCRs). GPCRs are Guanine nucleotide Exchange Factors (GEFs) that activate G-proteins by promoting the exchange of GDP for GTP on the Gα-subunits (Gilman, 1987). This signaling axis regulates a myriad of (patho)physiological processes and also represents the target for >30% of drugs on the market (Sriram and Insel, 2018), which fuels the interest in developing tools to manipulate G-protein activity in cells with high precision.

Optogenetics, the use of genetically-encoded proteins to control biological processes with light (Warden et al., 2014) is well-suited for the non-invasive manipulation of signaling. Metazoan opsins are light-activated GPCRs that have been repurposed as optogenetic tools (Airan et al., 2009; Bailes et al., 2012; Karunarathne et al., 2013; Oh et al., 2010; Siuda et al., 2015), albeit with limitations. For example, opsins tend to desensitize after activation due to receptor internalization and/or exhaustion of their chromophore, retinal (Airan et al., 2009; Bailes et al., 2012; Siuda et al., 2015). Exogenous supplementation of retinal, which is not always feasible, is required in many experimental settings because this chomophore is not readily synthesized by most mammalian cell types or by many organisms. Also, opsins are relatively large, post-translationally modified transmembrane proteins, which makes them inherently difficult to modify for optimization and/or customization for specific applications.

Here we leveraged the light-sensitive LOV2 domain of *Avena sativa* (Harper et al., 2003) to develop an optogenetic activator of heterotrimeric G-proteins not based on opsins. LOV2 uses ubiquitously abundant FMN as the chromophore and does not desensitize. It is small (∼17 KDa) and expresses easily as a soluble globular protein in different systems (Lungu et al., 2012; Strickland et al., 2012; Zayner et al., 2013), making it experimentally tractable, easily customizable, and widely applicable. Our results provide the proof of principle for a versatile optogenetic activator of heterotrimeric G-proteins that does not rely on GPCR-like proteins.

## RESULTS AND DISCUSSION

### Design and optimization of a photoswitchable G-protein activator

We envisioned the design of a genetically-encoded photoswitchable activator of heterotrimeric G-proteins by fusing a short sequence (∼25 amino acids) from the protein GIV (a.k.a. Girdin) called the *Gα-Binding-and-Activating* (GBA) motif to the C-terminus of LOV2 (**Fig. 1A, left**). GBA motifs are evolutionarily conserved sequences found in various cytoplasmic, non-GPCR proteins that are sufficient to activate heterotrimeric G-protein signaling (DiGiacomo et al., 2018). We reasoned that the GBA motif would be “uncaged” and bind G-proteins when the C-terminal Jα-helix of LOV2 becomes disordered upon illumination (Harper et al., 2003) (**Fig. 1A, right**). We named this optogenetic construct LOV2GIV (pronounced “*love-to-give”*). We used two well-validated mutants that mimic either the dark (D) or the lit (L) conformation of LOV2 (Harper et al., 2004; Harper et al., 2003; Zimmerman et al., 2016) to facilitate LOV2GIV characterization (**Fig. 1A, right**).

**Figure 1.**
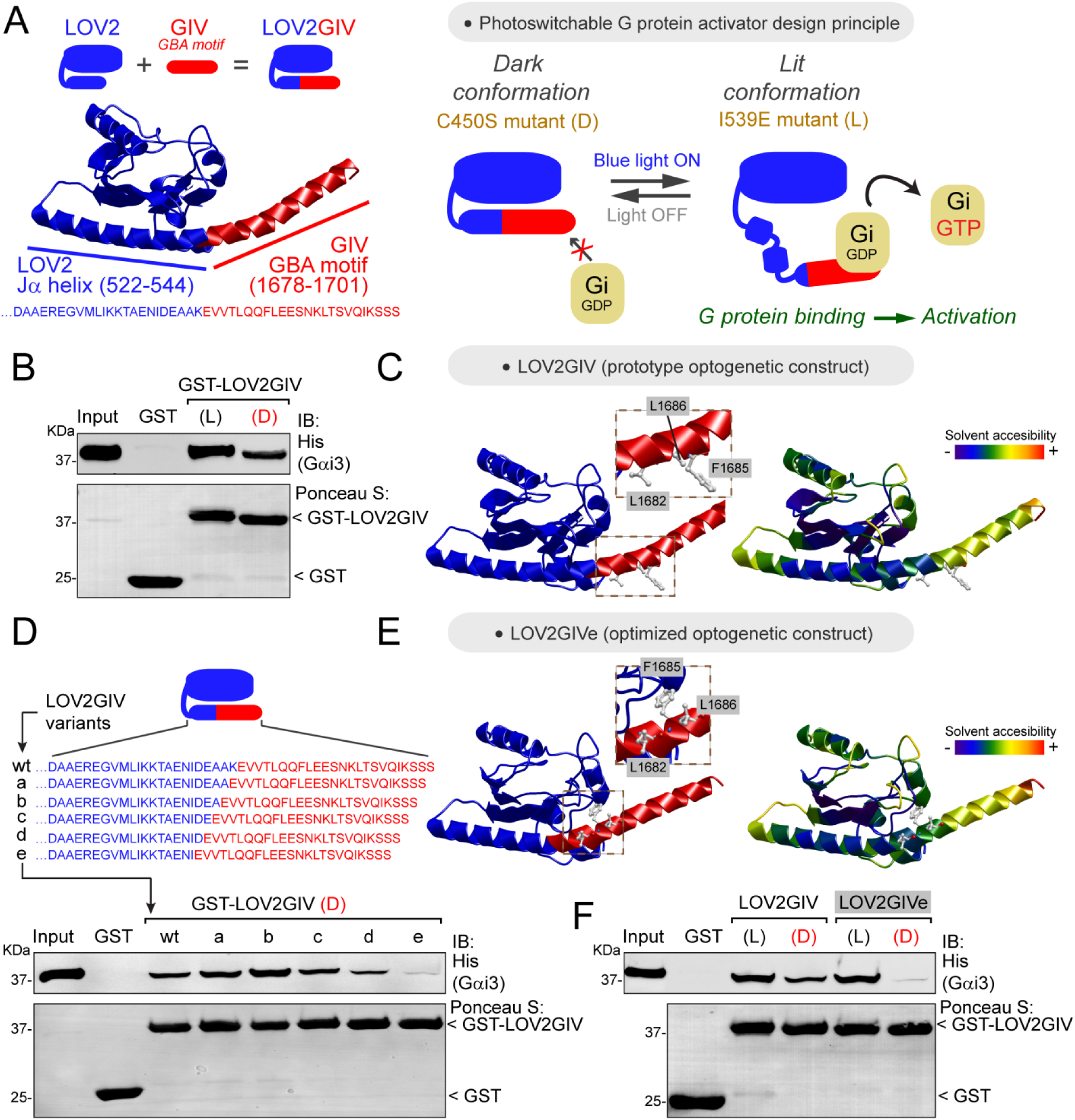
Design and optimization of LOV2GIV, an optogenetic activator of heterotrimeric G-proteins. **(A)** *<u>Left</u>*, diagram depicting the design of the LOV2GIV fusion protein. The GBA motif of GIV (red) is fused to the C-terminus of the LOV2 domain (blue). *<u>Right</u>*, diagram depicting the design principle of the photoswitchable G-protein activator LOV2GIV. In the dark conformation (D, which is mimicked by the LOV2 C450S mutant), the C-terminus of LOV2 forms an α-helix (Jα) and the GBA motif is not readily accessible for G-proteins. In the lit conformation (L, which is mimicked by the LOV2 mutant I539E), the C-terminus of LOV2 is more disordered and the GBA motif becomes accessible for G-proteins, which in turn are activated upon binding to the GBA motif. **(B)** LOV2GIV (L) binds Gαi3 better than LOV2GIV (D). Approximately 20 µg of the indicated purified GST-fused constructs were immobilized on glutathione-agarose beads and incubated with 3 µg (∼300 nM) of purified His-Gαi3. Resin-bound proteins were eluted, separated by SDS-PAGE, and analyzed by Ponceau S-staining and immunoblotting (IB) with the indicated antibodies. Input= 10% of the total amount of His-Gαi3 added in each binding reaction. **(C)** Ribbon representation of a LOV2GIV structure homology model generated using the I-TASSER server. On the left, the model is colored blue for LOV2 and red for GIV, whereas on the right it is colored according to solvent accessibility. Selected GIV residues known to be important for G-protein binding (L1682, F1685, L1686) (de Opakua et al., 2017; Kalogriopoulos et al., 2019) are displayed in stick representation and enlarged in the boxes. **(D)** LOV2GIV (D) variant “e” displays greatly diminished Gαi3 binding compared to LOV2GIV (D) wt. Protein binding experiments were carried out as described in panel B, except that the indicated LOV2GIV variants were used. **(E)** Ribbon representation of a structure homology model of the LOV2GIVe variant was generated and displayed as described for LOV2GIV in panel C. **(F)** The dynamic range of Gαi3 binding to lit versus dark conformations is improved for LOV2GIVe compared to LOV2GIV. Protein binding experiments were carried out as described in panel B, except that the indicated LOV2GIV variants were used. For all protein binding experiments in this figure, one representative result of at least three independent experiments is shown.

Our first LOV2GIV prototype had a suboptimal dynamic range, as it bound the G-protein Gαi3 in the lit conformation only ∼3 times more than in the dark conformation (**Fig. 1B**). Based on a structural homology model of LOV2GIV (**Fig. 1C**), we reasoned that it could be due to the relatively high accessibility of Gαi3-binding residues of the GIV GBA motif within the dark conformation of LOV2GIV. In agreement with this idea, fusion of the GBA motif at positions of the Jα helix more proximal to the core of the LOV2 domain tended to lower G-protein binding, including one variant (“e”, henceforth referred to as LOV2GIV<u>e</u>) in which binding to the dark conformation was almost undetectable (**Fig. 1D**). Consistent with this, a structure homology model of LOV2GIVe showed that amino acids required for G-protein binding are less accessible than in the LOV2GIV prototype (**Fig. 1E**). In contrast, the lit conformation of LOV2GIVe displayed robust Gαi3 binding, thereby confirming an improved dynamic range compared to the LOV2GIV prototype (**Fig. 1F**).

### LOV2GIVe binds and activates G-proteins efficiently *in vitro* only in its lit conformation

Concentration-dependent binding curves revealed that the affinity of Gαi3 for the dark conformation of LOV2GIVe is orders of magnitude weaker than for the lit conformation (**Fig. 2A**), which had an equilibrium dissociation constant (Kd) similar to that previously reported for the GIV-Gαi3 interaction (DiGiacomo et al., 2017). We also found that LOV2GIVe retains the same G-protein specificity as GIV. Much like GIV (Garcia-Marcos et al., 2010; Marivin et al., 2020), LOV2GIVe bound robustly only to members of the G_i/o_ family among the 4 families of Gα proteins (G_i/o_, G_s_, G_q/11_ and G_12/13_), and could discriminate within the G_i/o_ family by binding to Gαi1, Gαi2 and Gαi3 but not to Gαo (**Fig. 2B**). Next, we assessed if LOV2GIVe can activate G-proteins *in vitro*, i.e. whether it retains GIV’s GEF activity, by measuring steady-state GTPase activity as done previously for GIV and other related non-receptor GEFs (Garcia-Marcos et al., 2010; Maziarz et al., 2018). We found that LOV2GIVe in the lit, but not the dark, conformation led to a dose-dependent increase of Gαi3 activity (**Fig. 2C**), which was comparable to that previously shown for GIV (Garcia-Marcos et al., 2009). These findings indicate that LOV2GIVe recapitulates the G-protein activating properties of GIV *in vitro*, and that these are effectively suppressed for the dark conformation.

**Figure 2.**
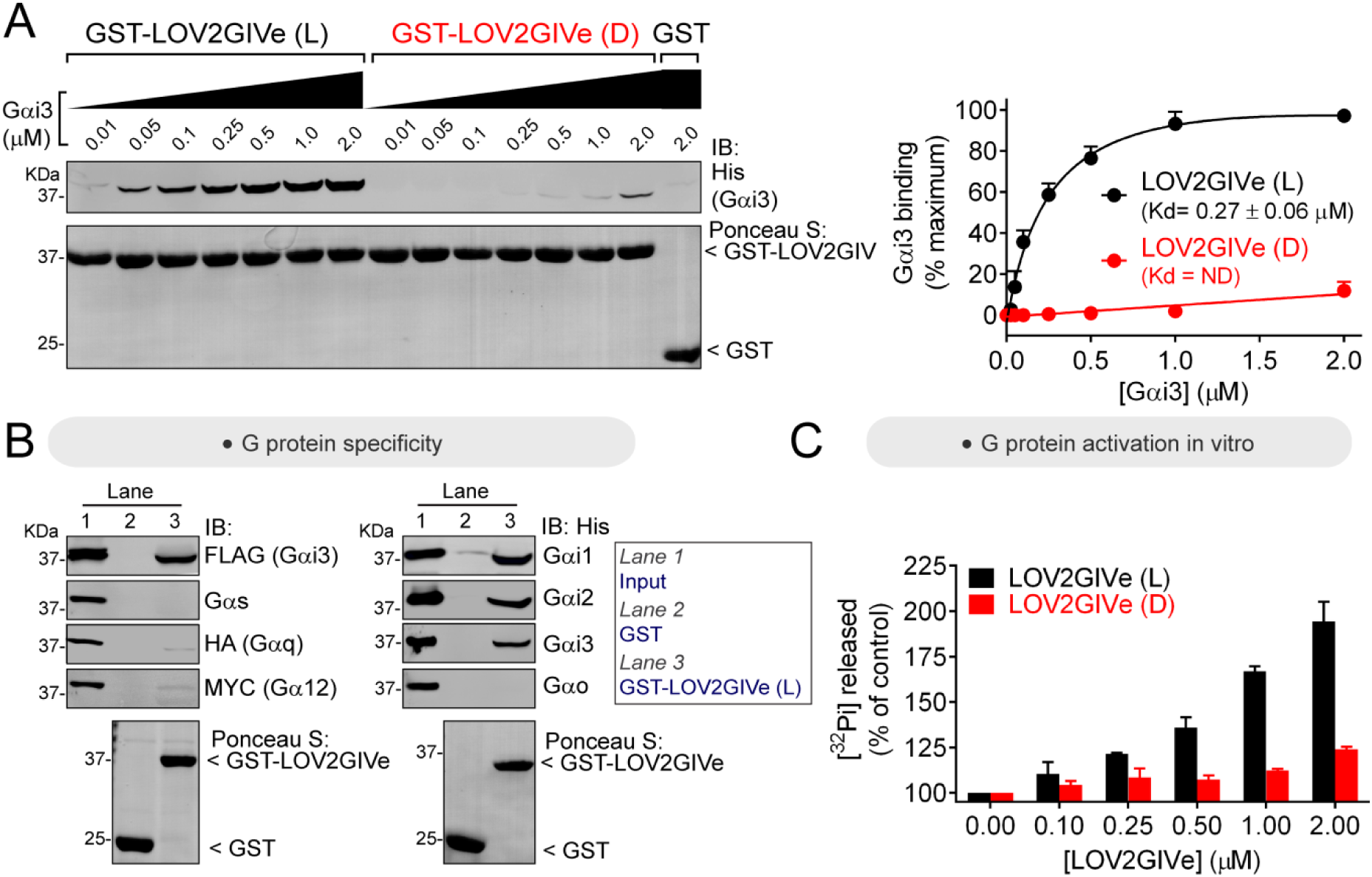
LOV2GIVe binds and activates Gαi3 *in vitro* in its lit conformation. **(A)** LOV2GIVe binds with high affinity to Gαi3. Approximately 10 µg of the indicated purified GST-fused constructs were immobilized on glutathione-agarose beads and incubated with the indicated concentrations of purified His-Gαi3. Resin-bound proteins were eluted, separated by SDS-PAGE, and analyzed by Ponceau S-staining and immunoblotting (IB) with the indicated antibodies. One representative result is shown on the left, and the graph on the right corresponds to the quantification of 3 independent experiments presented as mean ± S.E.M. for each data point and a solid line for the fit to a single site binding curve used to determine the Kd values. **(B)** LOV2GIVe binds specifically to Gαi compared to other Gα subunits. Approximately 20 µg of the indicated purified GST-fused constructs were immobilized on glutathione-agarose beads and incubated with the lysates of HEK293T cells expressing the indicated G-proteins (FLAG-Gαi3, Gαs, Gαq-HA and Gα12-MYC on the left panels) or purified His-tagged proteins (3 µg, ∼300 nM of His-Gαi1, His-Gαi2, His-Gαi3 and His-Gαo on the right panels). One representative experiment of at least three is shown. **(C)** LOV2GIVe (L), but not LOV2GIVe (D), increases Gαi3 activity *in vitro*. Steady-state GTPase activity of purified His-Gαi3 was determined in the presence of increasing amounts (0-2 µM) of purified GST-LOV2GIVe (L) (black) or GST-LOV2GIVe (D) (red) by measuring the production of [^32^P]Pi at 15 min. Results are the mean ± S.D. of n=2.

### LOV2GIVe activates G-protein signaling in cells in its lit conformation

To investigate LOV2GIVe-dependent G-protein activation in cells, we initially used a humanized yeast-based system (Cismowski et al., 1999; DiGiacomo et al., 2020) in which the mating pheromone response pathway has been co-opted to measure activation of human Gαi3 using a gene reporter (*LacZ*, β-galactosidase activity) (**Fig. 3A, left**). When we expressed LOV2GIVe dark (D), lit (L) or wt in this strain, only very weak levels of β-galactosidase activity were detected (**Fig. 3A, right**). We reasoned that this could be due to the subcellular localization of the construct, presumed to be cytosolic in the absence of any targeting sequence, because we have previously shown that GIV requires membrane localization to efficiently activate G-proteins (Parag-Sharma et al., 2016). Expressing LOV2GIVe (L) fused to a membrane targeting sequence (<u>m</u>LOV2GIVe) led to a very strong induction of β-galactosidase activity, which was not recapitulated by expression of mLOV2GIVe (D) or wt (**Fig. 3A, right**). LOV2GIVe-mediated activation levels (several hundred-fold over basal) are comparable to those previously reported for the endogenous pathway in response to GPCR activation by mating pheromone (Hoffman et al., 2002; Lambert et al., 2010). Next, we tested mLOV2GIVe in mammalian cells. Instead of using a downstream signaling readout as we did in yeast, we used a Bioluminescence Resonance Energy Transfer (BRET) biosensor that measures G-protein activity directly (**Fig. 3B, left**). Expression of mLOV2GIVe (L), but not (D), led to an increase of BRET proportional to the amount of plasmid transfected (**Fig. 3B, right**). At the highest amount tested, mLOV2GIVe (L) caused a BRET increase comparable to that observed upon agonist stimulation of the M4 muscarinic receptor, a Gi-activating GPCR. Introducing a mutation that precludes G-protein activation by GIV’s GBA motif (F1685A (FA), (Garcia-Marcos et al., 2009)), in mLOV2GIVe (L) also abolished its ability to induce a BRET increase (**Fig. 3-figure supplement 1**). These results indicate that the lit conformation of LOV2GIVe activates G-protein signaling in different cell types.

**Figure 3.**
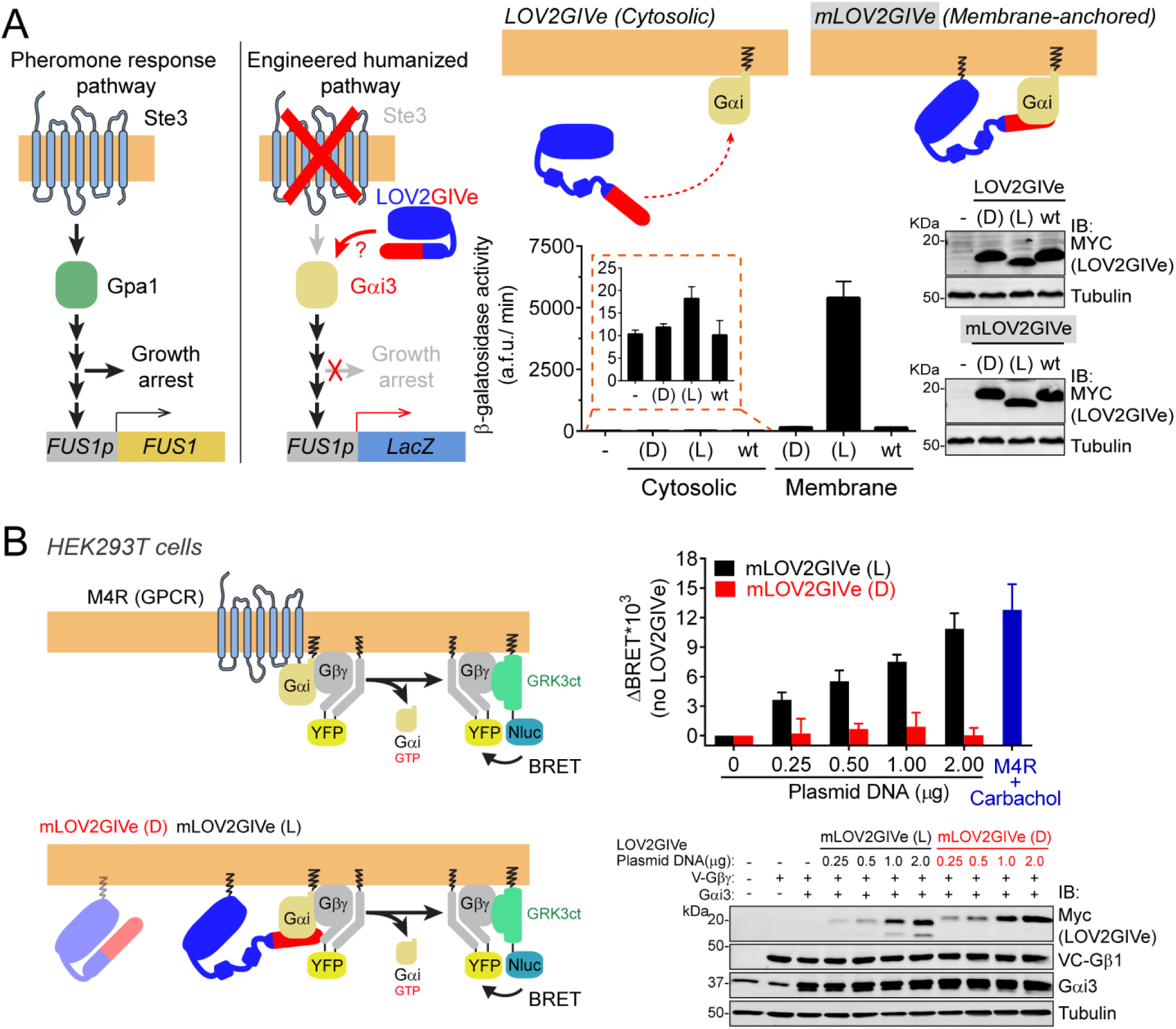
LOV2GIVe activates G-protein signaling in cells upon illumination. **(A)** *<u>Left</u>*, diagram comparing key steps and components of the mating pheromone response pathway of *S. cerevisiae* to those of an engineered, humanized strain used in the experiments shown in this figure. In the engineered strain, no pheromone responsive GPCR (Ste3) is expressed, the yeast G-protein Gpa1 is replaced by human Gαi3, and downstream signaling does not lead to growth arrest but promotes the activation of a reporter gene (*LacZ*) under the control of the pheromone-sensitive, G-protein-dependent promoter of *FUS1* (*FUS1p*). *<u>Right</u>*, membrane-anchored LOV2GIVe (mLOV2GIVe), but not its untargeted parental version, leads to strong G-protein activation only in the lit conformation. Yeast strains expressing the indicated LOV2GIVe constructs ((D), (L) or wt) or an empty vector (-) were lysed to determine β-galactosidase activity using a fluorogenic substrate (mean ± S.E.M., n=3) and to prepare samples for immunoblotting (one experiment representative of 3 is shown). **(B)** *<u>Left</u>*, diagrams showing the principle for the G-protein activity biosensor used in this panel. Upon action of a GPCR (top) or mLOV2GIVe (bottom), G-protein activation leads to the release of YFP-tagged Gβγ from Gαi, which then can associate with Nluc-tagged GRK3ct and results in the subsequent increase in BRET. *<u>Right</u>*, mLOV2GIVe (L) but not mLOV2GIVe (D), leads to increased G-protein activation similar to a GPCR as determined by BRET. BRET was measured in HEK293T cells transfected with the indicated amounts of mLOV2GIVe plasmid constructs along with the components of the BRET biosensor and the GPCR M4R. M4R was stimulated with 100 µM carbachol. Results in the graph on the top are expressed as difference in BRET (ΔBRET) relative to cells not expressing mLOV2GIVe (mean ± S.E.M., n=3). One representative immunoblot from cell lysates made after the BRET measurements is shown on the bottom.

### LOV2GIVe activates G-protein signaling in cells upon illumination

Next, we tested whether LOV2GIVe can trigger G-protein activation in cells in response to light. For this, we used two complementary experimental systems. The first one consisted of measuring G-protein activity directly with the mammalian cell BRET biosensor described above upon illumination with a pulse of blue light. We found that, in HEK293T cells expressing mLOV2GIVe wt, a single short light pulse resulted in a spike of G-protein activation that decayed at a rate similar to that reported for the transition from lit to dark conformation of LOV2 (T_1/2_ ∼1 min) (**Fig. 4A**). This response was not recapitulated by a mLOV2GIVe construct bearing the GEF-deficient FA mutation (**Fig. 4A**). For the second system, we turned to a spot growth assay with the humanized yeast stain described above. In this system, conditional histidine prototrophy controlled by *FUS1* promoter is used as a signaling readout downstream of G-protein activation under conditions of homogenous and continued illumination (**Fig. 4B, left**). Yeast cells expressing mLOV2GIVe wt grew in the absence of histidine only when exposed to blue light (**Fig. 4B, right**). This effect was specifically caused by light-dependent activation of mLOV2GIVe wt because cells expressing the light-insensitive mLOV2GIVe (D) construct failed to grow under the same illumination conditions, whereas mLOV2GIVe (L) grew the same regardless of illumination conditions (**Fig. 4B, right**). Taken together, these results show that mLOV2GIVe activates G-protein signaling in cells upon blue light illumination.

**Figure 4.**
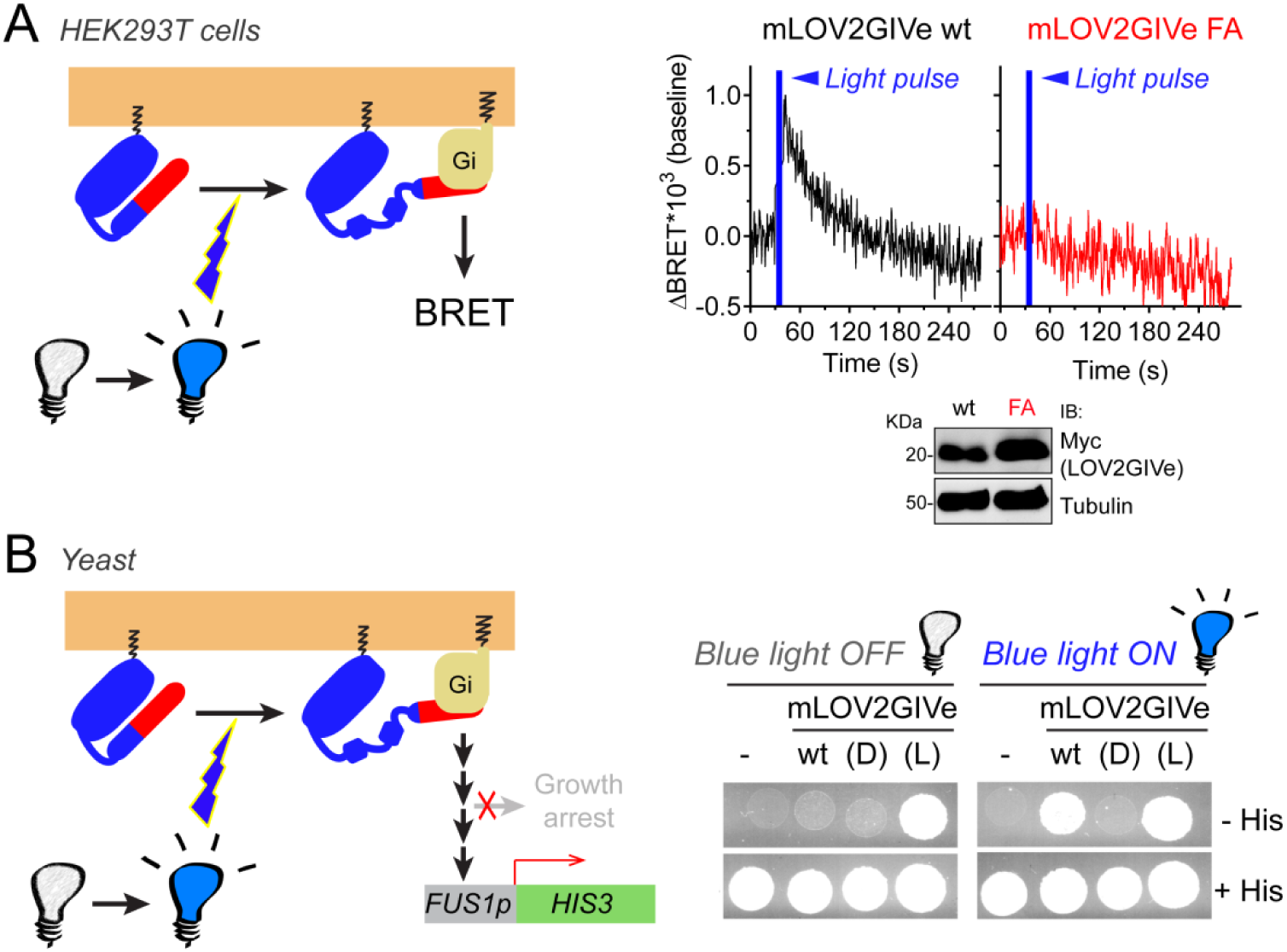
mLOV2GIVe activates G-protein signaling in cells upon illumination. **(A)** mLOV2GIVe activates G-proteins in mammalian cells upon illumination. Kinetic BRET measurements were carried out in HEK293T cells transfected with 2 µg of plasmids for the expression of mLOV2GIVe wt or FA along with the components of the BRET biosensor. Results are expressed as baseline corrected BRET changes (ΔBRET) and representative traces of one experiment out of four are presented. **(B)** mLOV2GIVe activates G-protein signaling in cells upon blue light illumination. Yeast strains expressing the indicated mLOV2GIVe constructs (wt, (D) or (L)) or an empty vector (-) were spotted on plates with or without histidine as indicated and imaged after 4 days of incubation in the dark (blue light OFF) or in the presence of blue light (blue light ON). One experiment representative of 3 is shown.

### Conclusions and future perspectives

Here we have presented proof-of-principle evidence for a photoswitchable G-protein activator that does not rely on opsins, i.e., light-activated GPCRs. This tool is based on a modular design that combines the properties of the light-sensitive LOV2 domain with a motif present in a non-GPCR activator of G-proteins. We propose that the versatility of the LOV2GIVe design could help overcome some limitations of currently available optogenetic tools that are based on GPCR-like proteins. For example, our results indicate that LOV2GIVe requires targeting to membranes where the substrate G-protein localizes (**Fig. 3A**), a feature that could be leveraged in the future to target it to different membranous organelles to study how activation of G-proteins in discrete subcellular compartments triggers different signaling events (Stoeber et al., 2018; Tsvetanova et al., 2015). Our results in yeast also support that LOV2GIVe can activate G-protein signaling (**Fig. 4C**) even in cell types where there is not sufficient synthesis of retinal to support opsin-based activation (Scott et al., 2019). A limitation of LOV2GIVe is that it only acts on one subset of heterotrimeric G-proteins, those containing Gαi subunits. Nevertheless, potential applications are still broad, as Gi proteins control processes as diverse as inhibitory neuromodulation, opioid action, or heart rate, among many others.

## MATERIALS AND METHODS

### Reagents and Antibodies

Unless otherwise indicated, all chemical reagents were obtained from Sigma or Fisher Scientific. Fluorescein di-β-D-galactopyranoside (FDG) was from Marker Gene Technologies, and the protein inhibitor mixture was from Sigma (catalog no. S8830). Leupeptin, pepstatin, and aprotinin were from Gold Biotechnology. All restriction endonucleases and *E. coli* strain BL21(DE3) were from Thermo Scientific. *E. coli* strain DH5α was purchased from New England Biolabs. *Pfu* ultra DNA polymerase used for site-directed mutagenesis was purchased from Agilent. [γ-^32^P]GTP was from Perkin Elmer. Mouse monoclonal antibodies raised against α-tubulin (T6074), FLAG tag (F1804) or His tag (H1029) were from Sigma. Mouse monoclonal antibody raised against hemagglutinin (HA) tag (clone 12CA5, #11583816001) was obtained from Roche. Mouse monoclonal antibody raised against MYC tag (9B11, #2276) was from Cell Signaling. Rabbit polyclonal antibody raised against Gαs (C-18, sc-383) was purchased from Santa Cruz Biotechnology. Goat anti-rabbit Alexa Fluor 680 (A21077) and goat anti-mouse IRDye 800 (#926-32210) secondary antibodies were from Life technologies and LiCor, respectively.

### Plasmid Constructs

The parental LOV2GIV sequence was obtained as a synthetic gene fragment from GeneScript and subsequently amplified by PCR with extensions at the 5’ and 3’ ends that made it compatible with a ligation independent cloning (LIC) system (Stols et al., 2002). For the bacterial expression of GST-LOV2GIV constructs, we inserted the LOV2GIV sequence into the pLIC-GST plasmid kindly provided by J. Sondek (UNC-Chapel Hill) (Cabrita et al., 2006). Two previously described LOV2 mutations (L514K and L531E) that reduce the spurious unwinding of the Jα helix in the dark conformation (Lungu et al., 2012; Zimmerman et al., 2016) were introduced in this construct. These and other mutations to generate the dark (D, C450S) and lit (L, I539E) conformations, the a, b, c, d, and e variants described on **Fig. 1D**, and the GEF-deficient FA mutant were made using the QuikChange II Mutagenesis Kit from Agilent. Full sequences of the parental LOV2GIV and LOV2GIVe variant are provided in **Supplementary Information**. For the yeast expression of LOV2GIVe constructs, we inserted the LOV2GIVe sequence into two different versions of a previously described pLIC-YES2 plasmid (Coleman et al., 2016). Both versions contain a MYC-tag sequence cloned upstream of the LIC cassette between the HindIII and KpnI sites, but in one of the two plasmids it was preceded by a sequence encoding the first 9 amino acids of *Saccharomyces cerevisiae* Gpa1, a previously validated membrane-targeting sequence (Parag-Sharma et al., 2016; Song et al., 1996). For the mammalian expression of LOV2GIVe constructs, we inserted the LOV2GIVe sequence into a modified pLIC-myc plasmid ((Cabrita et al., 2006), kindly provided by J. Sondek (UNC-Chapel Hill, NC)), in which a sequence encoding the first 11 amino acids of Lyn, a previously validated membrane-targeting sequence (Inoue et al., 2005; Parag-Sharma et al., 2016), was inserted in the AflII/KpnI sites upstream of the MYC-tag. The resulting sequence preceding the first amino acid of LOV2GIVe is **MGCIKSKGKD**SGTELGSM<u>EQKLISEEDL</u>GILYFQSNA (bold= Lyn11, underline= MYC-tag). Cloning of the pET28b-Gαi3 and pET28b-Gαo plasmids for the bacterial expression of rat His-Gαi3 or rat His-Gαo (isoform A), respectively, have been described previously (Garcia-Marcos et al., 2010; Garcia-Marcos et al., 2009). Plasmid pLIC-Gαi1(int.6xHis) for the bacterial expression of rat Gαi1 with an internal hexahistidine tag at position 120 was generated by PCR amplification from pQE6-Gαi1(int.6xHis) ((Dessauer et al., 1998), kindly provided by Carmen Dessauer, University of Texas, Houston) and insertion at NdeI/BglIIsites of the pLIC-His plasmid (Stols et al., 2002). The plasmid for the bacterial expression of rat His-Gαi2 (pET28b-Gαi2) was generated by inserting the Gαi2 sequence into the NdeI/EcoRI of pET28b. Plasmids for expression of FLAG-Gαi3 (rat, p3XFLAG-CMV10-Gαi3, N-terminal 3XFLAG tag) or Gαs (human, pcDNA3.1(+)-Gαs) in mammalian cells were described previously (Beas et al., 2012; Garcia-Marcos et al., 2009). Plasmids for the expression of Gαq-HA (mouse, pcDNA3-Gαq-HA, internally tagged) or Gα12-MYC (mouse, pcDNA3.1-Gα12-MYC, internally tagged) in mammalian cells were kindly provided by P. Wedegaertner (Thomas Jefferson University) (Wedegaertner et al., 1993) and T. Meigs (University of North Carolina, Asheville) (Ritchie et al., 2013), respectively. Plasmid for the expression of Gβ1 and Gγ2 fused to a split Venus (pcDNA3.1-Venus(1-155)-Gγ_2_ (VN-Gγ_2_) and pcDNA3.1-Venus(155-239)-Gβ_1_ (VC-Gβ_1_)) were kindly provided by N. Lambert (Augusta University, GA) (Hollins et al., 2009). pcDNA3.1-masGRK3ct-Nluc (Masuho et al., 2015) was a gift from K. Martemyanov (Scripps Research Institute, FL). The pcDNA3-Gαi3 plasmid for the expression of rat Gαi3 in mammalian cells has been described previously (Garcia-Marcos et al., 2010; Garcia-Marcos et al., 2009). The plasmid for the expression of M4R was obtained from the cDNA Resource Center at Bloomsburg University (pcDNA3.1-3xHA-M4R, cat# MAR040TN00).

### Protein Expression and Purification

All His-tagged and GST-tagged proteins were expressed in BL21(DE3) *E. coli* transformed with the corresponding plasmids by overnight induction at 23 °C with 1 mM isopropyl-β-D-1-thio-galactopyranoside (IPTG). Protein purification was carried out following previously described protocols (Garcia-Marcos et al., 2010; Garcia-Marcos et al., 2009). Briefly, bacteria pelleted from 1 L of culture were resuspended in 25 ml of buffer [50 mM NaH_2_PO_4_, pH 7.4, 300 mM NaCl, 10 mM imidazole, 1% (v:v) Triton X-100 supplemented with protease inhibitor cocktail (Leupeptin 1 µM, Pepstatin 2.5 µM, Aprotinin 0.2 µM, PMSF 1 mM)]. For His-Gαi3 and His-Gαo, this buffer was supplemented with 25 µM GDP and 5 mM MgCl_2_. After sonication (four cycles, with pulses lasting 20 s/cycle, and with 1 min interval between cycles to prevent heating), lysates were centrifuged at 12,000g for 20 min at 4 °C. The soluble fraction (supernatant) of the lysate was used for affinity purification on HisPur Cobalt or Glutathione Agarose resins (Pierce) and eluted with lysis buffer supplemented with 250 mM imidazole or with 50 mM Tris-HCl, pH 8, 100 mM NaCl, 30 mM reduced glutathione, respectively. GST-tagged proteins were dialyzed overnight at 4 °C against PBS. For His-Gαi1, His-Gαi2, His-Gαi3 and His-Gαo, the buffer was exchanged for 20 mM Tris-HCl, pH 7.4, 20 mM NaCl, 1 mM MgCl_2_, 1 mM DTT, 10 μM GDP, 5% (v/v) glycerol using a HiTrap Desalting column (GE Healthcare). All protein samples were aliquoted and stored at −80 °C.

### Protein-Protein Binding Assays

GST pulldown assays were carried out as described previously (Garcia-Marcos et al., 2010; Garcia-Marcos et al., 2011) with minor modifications. GST or GST-fused LOV2GIV constructs (described in “*Plasmid Constructs”*) were immobilized on glutathione agarose beads for 90 min at room temperature in PBS. Beads were washed twice with PBS, resuspended in 250-400 µl of binding buffer (50 mM Tris-HCl, pH 7.4, 100 mM NaCl, 0.4% (v:v) NP-40, 10 mM MgCl_2_, 5 mM EDTA, 1 mM DTT, 30 µM GDP) and incubated 4h at 4°C with constant tumbling in the presence of His-tagged G-proteins purified as described in “*Protein Expression and Purification*” or lysates of HEK293T cells expressing different G-proteins. For the latter, HEK293T cells (ATCC CRL-3216) were grown at 37°C, 5%CO_2_ in high-glucose Dulbecco’s Modified Eagle Medium (DMEM) supplemented with 10% FBS, 100U/ml penicillin, 100 µg/ml streptomycin and 1% L-glutamine. Approximately two million HEK293T cells were seeded on 10-cm dishes and transfected the day after using the calcium phosphate method with plasmids encoding the following constructs (DNA amounts in parenthesis): FLAG-Gαi3 (3 μg), Gαs (3 μg), Gαq-HA (6 μg) or Gα12-MYC (3 μg). Cell medium was changed 6 hours after transfection. Thirty two hours after transfection, cells were harvested by scraping in PBS and centrifugation before resuspension in 500 μl of ice-cold lysis buffer (20 mM Hepes, pH 7.2, 125 mM K(CH_3_COO), 0.4% (vol:vol) Triton X-100, 1 mM DTT, 10 mM β-glycerophosphate and 0.5 mM Na_3_VO_4_ supplemented with a protease inhibitor cocktail [SigmaFAST, #S8830]). Cell lysates were cleared by centrifugation at 14,000g for 10 min at 4°C. Approximately 20-25% of the lysate from a 10-cm dish was used for each binding reaction. After incubation with purified proteins or cell lysates, beads were washed four times with 1 ml of wash buffer (4.3 mM Na_2_HPO_4_, 1.4 mM KH_2_PO_4_, pH 7.4, 137 mM NaCl, 2.7 mM KCl, 0.1% (v/v) Tween-20, 10 mM MgCl_2_, 5 mM EDTA, 1 mM DTT and 30 µM GDP) and resin-bound proteins eluted by boiling for 5 min in Laemmli sample buffer before processing for immunoblotting (see below “*Immunoblotting*”).

### Protein Structure Modeling and Visualization

Models of LOV2GIV and LOV2GIVe were generated using the server I-TASSER (https://zhanglab.ccmb.med.umich.edu/I-TASSER/, (Yang et al., 2015)). Best scoring models were chosen for further analysis (LOV2GIV C-score= −0.64, LOV2GIVe C-score= −0.33). Protein structures were visualized and displayed using ICM version 3.8-3 (Molsoft LLC., San Diego, CA).

### Steady-state GTPase Assay

This assay was performed as described previously (Garcia-Marcos et al., 2010; Garcia-Marcos et al., 2009; Garcia-Marcos et al., 2011). Briefly, His-Gαi3 (400 nM) was preincubated with different concentrations of GST-LOV2GIVe constructs for 15 min at 30°C in assay buffer [20 mM Na-HEPES, pH 8, 100 mM NaCl, 1 mM EDTA, 25 mM MgCl_2_, 1 mM DTT, 0.05% (w:v) C_12_E_10_]. GTPase reactions were initiated at 30°C by adding an equal volume of assay buffer containing 1 µM [γ-^32^P]GTP (∼50 c.p.m/ fmol). Duplicate aliquots (25 μl) were removed at 15 min and reactions stopped with 975 μl of ice-cold 5% (w/v) activated charcoal in 20 mM H_3_PO_4_, pH 3. Samples were then centrifuged for 10 min at 10,000 g, and 500 μl of the resultant supernatant were scintillation counted to quantify [^32^P]Pi released. Background [^32^P]Pi detected at 15 min in the absence of G-protein was subtracted from each reaction and data expressed as percentage of the Pi produced by His-Gαi3 in the absence of GST-LOV2GIVe. Background counts were <5% of the counts detected in the presence of G-proteins.

### Yeast Strains and Manipulations

The previously described (Cismowski et al., 1999) *Saccharomyces cerevisiae* strain CY7967 [*MATα GPA1(1-41)-Gαi3 far1*Δ *fus1p-HIS3 can1 ste14:trp1:LYS2 ste3*Δ *lys2 ura3 leu2 trp1 his3*] (kindly provided by James Broach, Penn State University) was used for all yeast experiments. The main features of this strain are that the gene encoding only pheromone responsive GPCR (*STE3*) is deleted, the endogenous Gα-subunit Gpa1 is replaced by a chimeric Gpa1(1-41)-human Gαi3 (36-354) and the gene encoding the cell cycle arrest-inducing protein Far1 is deleted. In this strain, the pheromone response pathway can be upregulated by the ectopic expression of activators of human Gαi3 and does not result in the cell cycle arrest that occurs in the native pheromone response (Cismowski et al., 2002; Cismowski et al., 1999; Maziarz et al., 2018). Plasmid transformations were carried out using the lithium acetate method. CY7967 was first transformed with a centromeric plasmid (CEN TRP) encoding the *LacZ* gene under the control of the *FUS1* promoter (*FUS1p*), which is activated by the pheromone response pathway. The *FUS1p*::*LacZ*-expressing strain was transformed with pLIC-YES2 plasmids (2µm, URA) encoding each of the LOV2GIV constructs described in “*Plasmid Constructs”*. Double transformants were selected in synthetic defined (SD)-TRP-URA media. Individual colonies were inoculated into 3 ml of SDGalactose-TRP-URA and incubated overnight at 30°C to induce the expression of the proteins of interest under the control of the galactose-inducible promoter of pLIC-YES2. This starting culture was used to inoculate 20 ml of SDGalactose-TRP-URA at 0.3 OD600. Exponentially growing cells (∼0.7-0.8 OD600, 4-5 hours) were pelleted to prepare samples for “*β-galactosidase Activity Assay*” and “*Yeast Spot Growth Assay*” described below and for preparing samples for immunobloting as previously described (de Opakua et al., 2017; Maziarz et al., 2018). Briefly, pellets corresponding to 5 OD600 were washed once with PBS + 0.1% BSA and resuspended in 150 µl of lysis buffer (10 mM Tris-HCl, pH 8.0, 10% (w:v) trichloroacetic acid (TCA), 25 mM NH_4_OAc, 1 mM EDTA). 100 µl of glass beads were added to each tube and vortexed at 4°C for 5 min. Lysates were separated from glass beads by poking a hole in the bottom of the tubes followed by centrifugation onto a new set of tubes. The process was repeated after the addition of 50 µl of lysis buffer to wash the glass beads. Proteins were precipitated by centrifugation (10 min, 20,000 g) and resuspended in 60 µl of solubilization buffer (0.1 M Tris-HCl, pH 11.0, 3% SDS). Samples were boiled for 5 min, centrifuged (1 min, 20,000 g) and 50 µl of the supernatant transferred to new tubes containing 12.5 µl of Laemmli sample buffer and boiled for 5 min.

### β-galactosidase Activity Assay

This assay was performed as described previously (Hoffman et al., 2002; Maziarz et al., 2018) with minor modifications. Pellets corresponding to 0.5 OD600 (in duplicates) were washed once with PBS + 0.1% (w:v) BSA and resuspended in 200 µl assay buffer (60 mM Na_2_PO_4_, 40 mM NaH_2_PO_4_, 10 mM KCl, 1 mM MgCl_2_, 0.25% (v:v) β-mercaptoethanol, 0.01% (w:v) SDS, 10% (v:v) chloroform) and vortexed. 100 µl were transferred to 96-well plates and reactions started by the addition of 50 µl of the fluorogenic β-galactosidase substrate fluorescein di-β-D-galactopyranoside (FDG, 100 µM final). Fluorescence (Ex. 485 ± 10 nm/ Em. 528 ± 10 nm) was measured every 2 min for 90 min at 30°C in a Biotek H1 synergy plate reader. Enzymatic activity was calculated as arbitrary fluorescent units (a.f.u.) per minute (min).

### Bioluminescence Resonance Energy Transfer (BRET)-based G Protein Activation Assay

BRET experiments were conducted as described previously (Maziarz et al., 2018). HEK293T cells (ATCC®, CRL-3216) were seeded on 6-well plates (∼400,000 cells/well) coated with gelatin and after one day transfected using the calcium phosphate method with plasmids encoding for the following constructs (DNA amounts in parenthesis): Venus(155-239)-Gβ_1_ (VC-Gβ_1_) (0.2 μg), Venus(1-155)-Gγ_2_ (VN-Gγ_2_) (0.2 μg) and Gαi3 (1 μg) mas-GRK3ct-Nluc (0.2 μg) along with mLOV2GIVe constructs in the amounts indicated in the corresponding figure legends. Approximately 16-24 hours after transfection, cells were washed and gently scraped in room temperature PBS, centrifuged (5 min at 550g) and resuspended in assay buffer (140 mM NaCl, 5 mM KCl, 1 mM MgCl_2_, 1 mM CaCl_2_, 0.37 mM NaH_2_PO_4_, 20 mM HEPES pH 7.4, 0.1% glucose) at a concentration of 1 million cells/ml. 25,000-50,000 cells were added to a white opaque 96-well plate (Opti-Plate, Perkin Elmer) and mixed with the nanoluciferase substrate Nano-Glo (Promega cat# N1120, final dilution 1:200) for 2 min before measuring luminescence signals in a POLARstar OMEGA plate reader (BMG Labtech) at 28 °C. Luminescence was measured at 460 ± 40 nm and 535 ± 10 nm, and BRET was calculated as the ratio between the emission intensity at 535 ± 10 nm divided by the emission intensity at 460 ± 40 nm. For the activation of M4R in **Fig. 3B**, cells were exposed to 100 µM carbachol for 4 min prior to measuring BRET. For measurements shown in **Fig. 3B** and **Fig. 4B**, BRET data are presented as the difference from cells not expressing LOV2GIVe constructs. For kinetic BRET measurements shown in **Fig. 4A**, luminescence signals were measured every 0.24 seconds for the duration of the experiment. The illumination pulse was achieved by switching from the luminescence read mode to the fluorescence read mode of the plate reader, with the following settings: 485 ± 6 nm filter, 200 flashes (∼1.5 s). After the pulse of illumination, measurements were returned to luminescence mode with the same settings as prior to illumination. BRET data was corrected by subtracting the BRET signal baseline (average of 30 seconds pre-light pulse) and then subjected to a smoothening function (second order, four neighbors) in GraphPad for presentation. At the end of some BRET experiments, a separate aliquot of the same pool of cells used for the luminescence measurements was centrifuged for 1 min at 14,000 g and pellets stored at −20 °C. To prepare lysates for immunoblotting, pellets were resuspended in lysis buffer (20 mM HEPES, pH 7.2, 5 mM Mg(CH3COO)_2_, 125 mM K(CH3COO), 0.4% Triton X-100, 1 mM DTT, and protease inhibitor mixture). After clearing by centrifugation at 14,000 g at 4 °C for 10 min, protein concentration was determined by Bradford and samples boiled in Laemmli sample buffer for 5 min before following the procedures described in “*Immunoblotting*”.

### Yeast Spot Growth Assay

Cells bearing LOV2GIVe constructs growing exponentially in SDGalactose media were pelleted as described above (“*Yeast Strains and Manipulations”*), and resuspended at equal densities. Equal volumes of each strain were spotted on agar plates in 4 identical sets. Two of the sets were seeded on SDGalactose-TRP-URA plates with histidine and the other two sets were seeded on SDGalactose-TRP-URA-HIS (supplemented with 5 mM 3-amino-1,2,4-triazole). From each one of the two pairs of sets, one of the plates was exposed to a homemade array of blue LED strips positioned approximately 12 cm above the plates (∼2,000 Lux as determined by a Trendbox Digital Light Meter HS1010A) whereas the other one was incubated side by side under the same light but tightly wrapped in aluminum foil. Plates were incubated simultaneously under these conditions for 4 days at 30 °C and then imaged using an Epson flatbed scanner.

### Immunoblotting

Proteins were separated by SDS-PAGE and transferred to PVDF membranes, which were blocked with 5% (w:vol) non-fat dry milk and sequentially incubated with primary and secondary antibodies. For protein-protein binding experiments with GST-fused proteins, PVDF membranes were stained with Ponceau S and scanned before blocking. The primary antibodies used were the following: MYC (1:1,000), His (1:2,500), FLAG (1:2,000), α-tubulin (1:2,500), HA (1:1,000) and Gαs (1:500). The secondary antibodies were goat anti-rabbit Alexa Fluor 680 (1:10,000) and goat anti-mouse IRDye 800 (1:10,000). Infrared imaging of immunoblots was performed using an Odyssey Infrared Imaging System (Li-Cor Biosciences). Images were processed using the ImageJ software (NIH) and assembled for presentation using Photoshop and Illustrator softwares (Adobe).

## AUTHOR CONTRIBUTIONS

MG-M, KP-S, AM, MM and LTN conducted experiments. AL provided critical reagents. MG-M conceived the project, designed experiments, supervised experiments and wrote the manuscript.

## ACKNOWLEDGEMENTS

This work was supported by NIH grant R01GM136132 (to MG-M). MM was supported by an American Cancer Society-Funding Hope Postdoctoral Fellowship, PF-19-084-01-CDD. We thank N. Lambert (Augusta University), K. Martemyanov (The Scripps Research Institute, Florida), C. Dessauer (University of Texas, Houston), J. Sondek (University of North Carolina-Chapel Hill), P. Wedegaertner (Thomas Jefferson University) and T. Meigs (University of North Carolina-Asheville) for providing plasmids.

---

**Figure 3- figure supplement 1.**
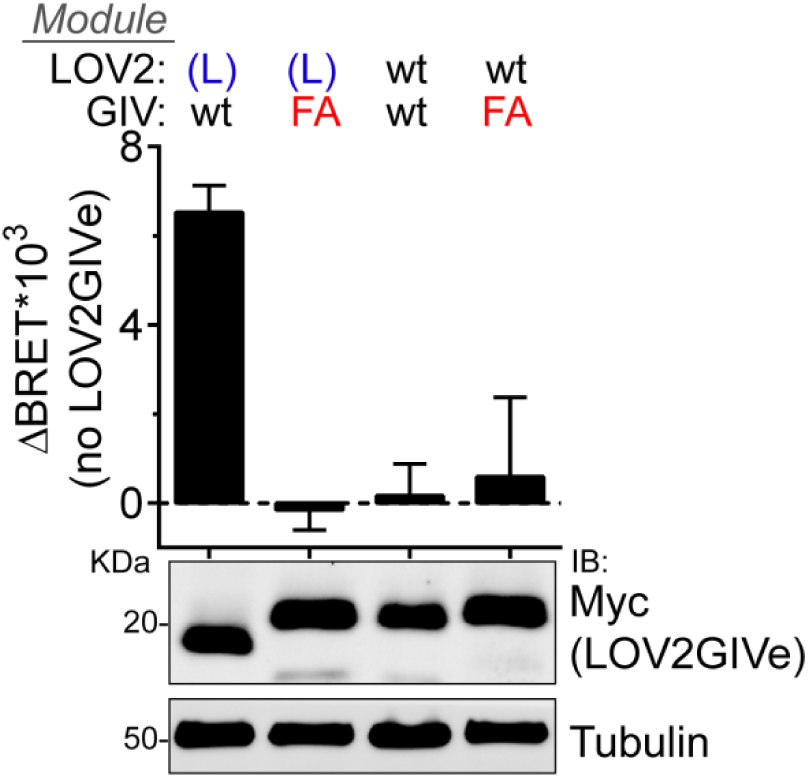
FA mutation abolishes G-protein activation by mLOV2GIVe (L). BRET was measured in HEK293T cells transfected with 2 µg of the indicated mLOV2GIVe plasmid constructs along with the components of the BRET biosensor described in **Fig. 3**. Results in the graph on the top are expressed as difference in BRET (ΔBRET) relative to cells not expressing mLOV2GIVe (mean ± S.E.M., n=5). One representative immunoblot from cell lysates made after the BRET measurements is shown on the bottom.

## SUPPLEMENTARY INFORMATION

Sequences of LOV2GIV constructs

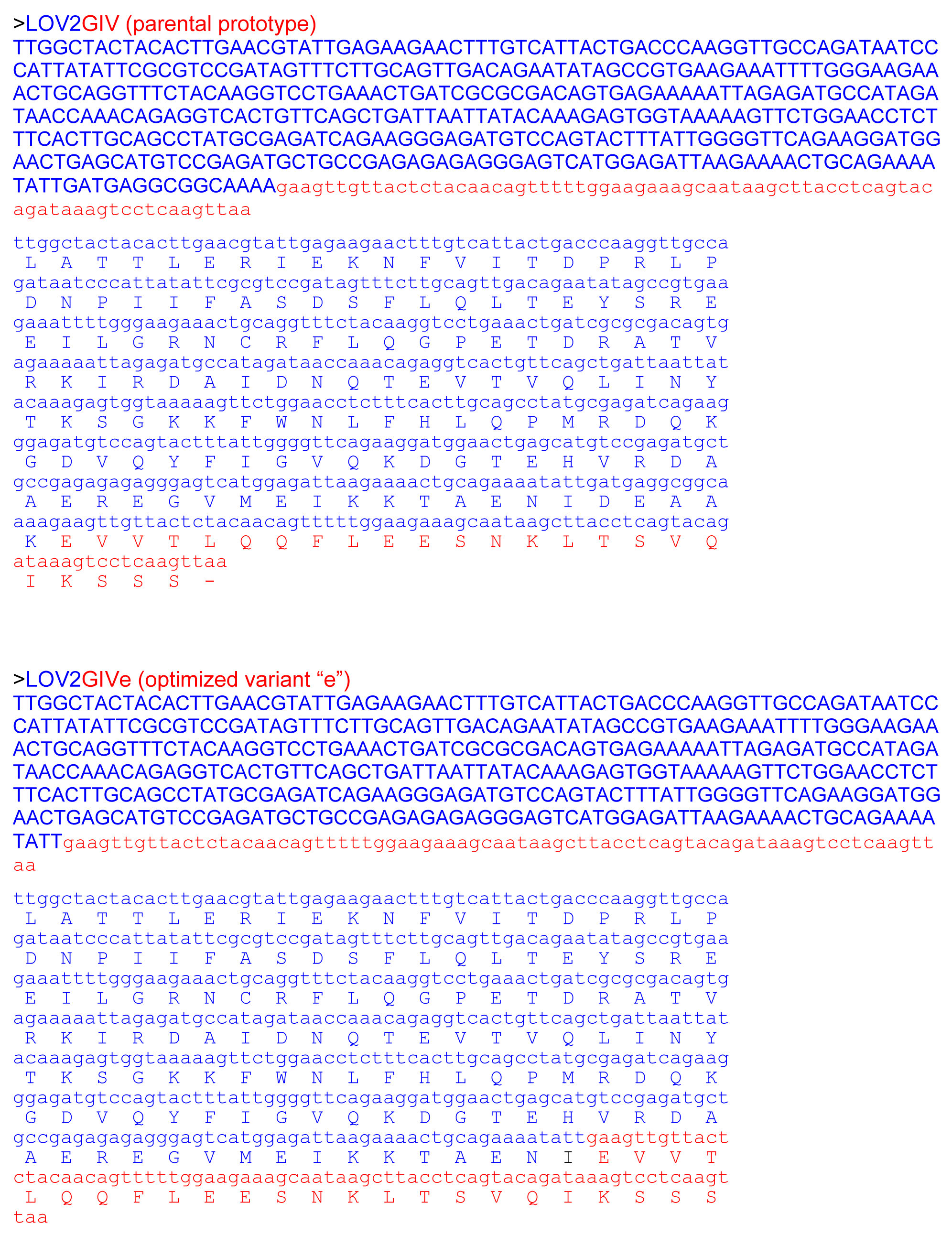

